# *In situ* architecture of the endosymbiont *Wolbachia pipientis*

**DOI:** 10.1101/2025.08.29.673095

**Authors:** Sujit Pujhari, Jessica Heebner, Erich Raumann, Tengfei Zhong, Jason L. Rasgon, Matthew T. Swulius, Carrie L. Shaffer, Mohammed Kaplan

## Abstract

Hidden within host cells, the endosymbiont *Wolbachia pipientis* is the most prevalent bacterial infection in the animal kingdom. Scientific breakthroughs over the past century yielded fundamental mechanisms by which *Wolbachia* controls arthropod reproduction to shape dynamic ecological and evolutionary trajectories. However, the structure and spatial organization of symbiont machineries that underpin intracellular colonization and orchestrate maternal inheritance remain unknown. Here, we used cryo-electron tomography to directly image the nanoscale architecture of bacterial tools deployed for host manipulation and germline transmission. We discovered that *Wolbachia* assembles multiple structures at the host-endosymbiont interface including a filamentous ladder-like framework hypothesized to serve as a specialized motility mechanism that enables bacterial translocation to specific host cell compartments during embryogenesis and somatic tissue dissemination. In addition, we present the first *in situ* structure of the Rickettsiales *vir* homolog type IV secretion system (*rvh* T4SS). We provide evidence that the *rvh* T4SS nanomachine exhibits architectural similarities to the pED208-encoded T4SS apparatus including the biogenesis of rigid conjugative pili extending hundreds of nanometers beyond the bacterial cell surface. Coupled with integrative structural modeling, we demonstrate that in contrast to canonical T4SS architectures, the α-proteobacterial T4SS outer membrane complex assembles a periplasmic baseplate structure predicted to comprise VirB9 oligomers complexed with cognate VirB10 subunits that form extended antennae projections surrounding the translocation channel pore. Collectively, these studies provide an unprecedented view into *Wolbachia* structural cell biology and unveil the molecular blueprints for architectural paradigms that reinforce ancient host-microbe symbioses.

## INTRODUCTION

Microbial endosymbiosis plays a central role in eukaryotic cell origin, speciation, ecological interactions, animal behavior, invertebrate reproduction, sex determination, and embryogenesis (*1–3*). Arguably the most significant and evolutionarily remarkable endosymbiotic microbe, the Gram-negative, obligate intracellular bacterium *Wolbachia pipientis* (“*Wolbachia*”) is harbored by diverse invertebrates belonging to the Ecdysozoa superphylum (*4*). Infecting approximately 40-50% of terrestrial arthropods and several onchocercid nematode species worldwide (*1–3*), *Wolbachia* exhibits either mutualistic or parasitic mechanisms depending on the host species. In arthropods, *Wolbachia* imparts fitness advantages for infected females and enables maternal germ line bacterial inheritance by manipulating host reproduction via male killing (*1, 5, 6*), feminization (*7*), parthenogenesis (*8*), and cytoplasmic incompatibility (CI) mechanisms (*1, 9*). In some vector arthropod systems, the bacterium reduces vectorial capacity by providing immunity to viral and parasitic infections including dengue virus, Zika virus, Chikungunya virus, yellow fever virus, O’nyong-nyong virus, Mayaro virus, and malaria (*10–19*), and enhances reproductive fitness by modulating host iron homeostasis (*20*). *Wolbachia* fulfills a central obligatory mutualistic role in onchocercid nematodes, whereby infection fosters embryogenesis, development, and filariae survival (*3, 21, 22*); in exchange, the bacterium obtains essential amino acids, vitamins, and cofactors from the host, demonstrating complex interkingdom metabolic dependencies (*23, 24*). Together with the ability of the bacterium to switch hosts and transmit vertically, harnessing *Wolbachia* endosymbiosis thus represents a promising biocontrol strategy to reduce devastating arbovirus and filarial infections in the developing world. For example, *Wolbachia* spread (mediated by CI) has emerged as an effective and sustainable *Aedes aegypti* vector control strategy that simultaneously decreases pathogen replication and transmission (*15, 16, 25–27*). Similarly, therapeutically targeting *Wolbachia* mutualists in filarial nematodes demonstrates the impact of endosymbiont-based strategies to eliminate diseases such as lymphatic filariasis and river blindness that together threaten over one billion people globally (*28–30*).

Within both arthropod and nematode hosts, *Wolbachia* colonize diverse reproductive and somatic tissues (*30, 31*). In *Drosophila* ovaries, *Wolbachia* are localized in the vicinity of various organelles and structures including the endoplasmic reticulum (ER) (*32*), Golgi apparatus (*33*), and the actin cytoskeleton (*34*) and microtubule network (*35, 36*) which are exploited for bacterial intracellular trafficking and tissue colonization during germline cell differentiation. Previous studies demonstrate that while *Wolbachia* exhibits prototypical Gram-negative inner and outer membranes, an additional third membrane, presumably derived from localized ER remodeling, surrounds the endosymbiont (*30, 37*), resulting in altered host cell organization and vesicle trafficking pathway disruption (*38, 39*). Interestingly, *Wolbachia* exhibits various morphologies within the host cell cytoplasm including spherical, rod, and intermediate forms (*37, 40–43*); however, the functional significance of *Wolbachia* pleomorphism is unresolved.

Recent dual host-microbe sequencing efforts have shed light on potential mechanisms by which *Wolbachia* dynamically interacts with the protected intracellular niche during tissue colonization and transmission (*24, 44, 45*). For example, transcriptional analysis across the *Drosophila* lifespan revealed that *Wolbachia* synthesize components of a Rickettsiales *vir* homolog type IV secretion system (*rvh* T4SS) throughout host development from embryogenesis to adulthood in both male and female flies (*45–47*). While functional characterization of putative effector proteins has been hampered by a lack of conventional molecular tools and *Wolbachia* genetic intractability, exciting studies leveraging heterologous systems (*48*), surrogate eukaryotic models (*49–52*), and *Drosophila* genetics (*49, 53, 54*) have uncovered novel candidate substrates that are presumed to be translocated into host cells via *w*Mel *rvh* T4SS-dependent mechanisms. Previous studies focused on genomic architecture and evolution suggest that while the Rickettsiales T4SS machinery evolved from paradigmatic bacterial conjugation systems (*55, 56*), α-proteobacterial architectures incorporate specialized modifications tailored to facilitate the endosymbiont lifestyle.

For instance, in comparison to the prototypical *Agrobacterium tumefaciens vir* T4SS, the *Wolbachia w*Mel *rvh* T4SS apparatus lacks canonical VirB1 (lytic transglycosylase), VirB5 (minor subunit of the conjugative pilus), and VirB7 (outer membrane complex component) orthologs (*46, 47, 57*). As with other Rickettsiales, *Wolbachia* encodes pseudogenized or truncated T4SS components as well as multiple, presumably functionally redundant copies of *virB6* (secretion channel/stalk protein), *virB9* (outer membrane complex component), *virB8* (inner membrane component), *virB4* (ATPase), and *virB2* (conjugative pilin) orthologs (*46, 57*). The observation that some machinery components and putative T4SS effectors are differentially regulated (i) during early larval and mid-to-late pupal stages of *D. melanogaster* development and (ii) in a male sex-dependent manner in adult flies (*45, 46*) suggests that *Wolbachia* exploits *rvh* T4SS mechanisms to manipulate host biology in response to specific host developmental cues. Notably, multiple *virB2* loci exhibit striking upregulation during pupation, suggesting an important role of the putative *rvh* T4SS conjugative pilus in host manipulation. However, whether an external pilus is assembled or required for *w*Mel *rvh* T4SS effector and/or DNA transport into either host cells or co-resident *Wolbachia* is unknown.

Previous studies of *Wolbachia* ultrastructure relied mainly on light microscopy and conventional transmission electron microscopy (TEM) techniques to visualize the host-endosymbiont interface (*32, 33, 35, 37, 40, 41, 43, 58, 59*). However, several experimental limitations associated with sample preparation methods confound data interpretation. For example, sample fixation, dehydration, and staining techniques employed in conventional TEM disrupt membranes and result in artefacts that obscure macromolecular details, thereby reducing ultrastructural resolution. To overcome these experimental challenges, we used cryo-electron tomography (cryo-ET) to directly visualize isolated *Wolbachia w*Mel in the native, frozen-hydrated state at macromolecular resolution (*60, 61*). We discovered several previously uncharacterized cellular features including periplasmic and inner membrane-associated complexes, ladder-like structures assembled on the *Wolbachia* cell surface, and multiple architectural states of the *w*Mel *rvh* T4SS nanomachine. Notably, we captured the biogenesis of rigid, extracellular conjugative pili associated with the assembled *w*Mel *rvh* T4SS apparatus. Coupled to the identification of other novel macromolecular structures that open new research avenues, our studies provide an unprecedented view into enigmatic *Wolbachia* cell biology and highlight the fascinating complexity of microbial machinery assembled at the host-endosymbiont interface.

## RESULTS

### *Wolbachia* exhibits diverse morphologies during host colonization

Previous studies employing TEM methodologies reported spherical, rod shaped, and intermediate *Wolbachia* morphologies identified within cognate arthropod and nematode host tissues (*37, 40–43*). However, detailed ultrastructural analysis has not been performed in the context of isolated *Wolbachia* cells liberated from the host. To gain insight into structural mechanisms underlying the endosymbiont lifestyle, we purified *Wolbachia w*Mel from cultured *Anopheles* mosquito cells and imaged isolated bacteria in the native state using cryo-ET. In accordance with previous studies (*37, 40, 41, 43*), purified *Wolbachia* exhibited various developmental morphologies ranging from bacillus (28%) to spherical (36%) and intermediate coccobacillus forms (36%) (**Fig. 1A-C** and **Table S1**). Consistent with canonical Gram-negative architecture, isolated *Wolbachia* exhibited a characteristic outer membrane (OM) and inner membrane (IM) (**Fig. 1**). Interestingly, the *Wolbachia* OM presented as either smooth (23 cells, **Fig. 1A**) or ruffled (18 cells, **Fig. 1C**) forms with no clear correlation observed between cell morphology (either spheroid or bacillus) and the nature of the OM. We additionally identified examples of *Wolbachia* undergoing symmetrical cell division (**Fig. 1D**). Isolated cells were frequently observed in clusters (**Fig. 1E-F**) reminiscent of *Wolbachia* aggregates localized within host cell inclusion vacuoles previously imaged by conventional TEM (*43*). Resulting from sample preparation relics, we also observed rare instances of isolated *Wolbachia* associated with purified *Anopheles-*derived cellular membranes (**Fig. 1G-I** and **Movie S1**). However, analysis of individual tomograms and segmentation (**Movie S1**) did not reveal either loose association or direct contact between the *Wolbachia* OM and the surrounding host cell membrane (**Fig. 1G-I** and **Movie S1**). Collectively, these results corroborate and build upon previous TEM observations of *Wolbachia* imaged in the context of host cell colonization and provide the first ultrastructural visualization of isolated endosymbiont cells in a near-native state.

**Figure 1.**
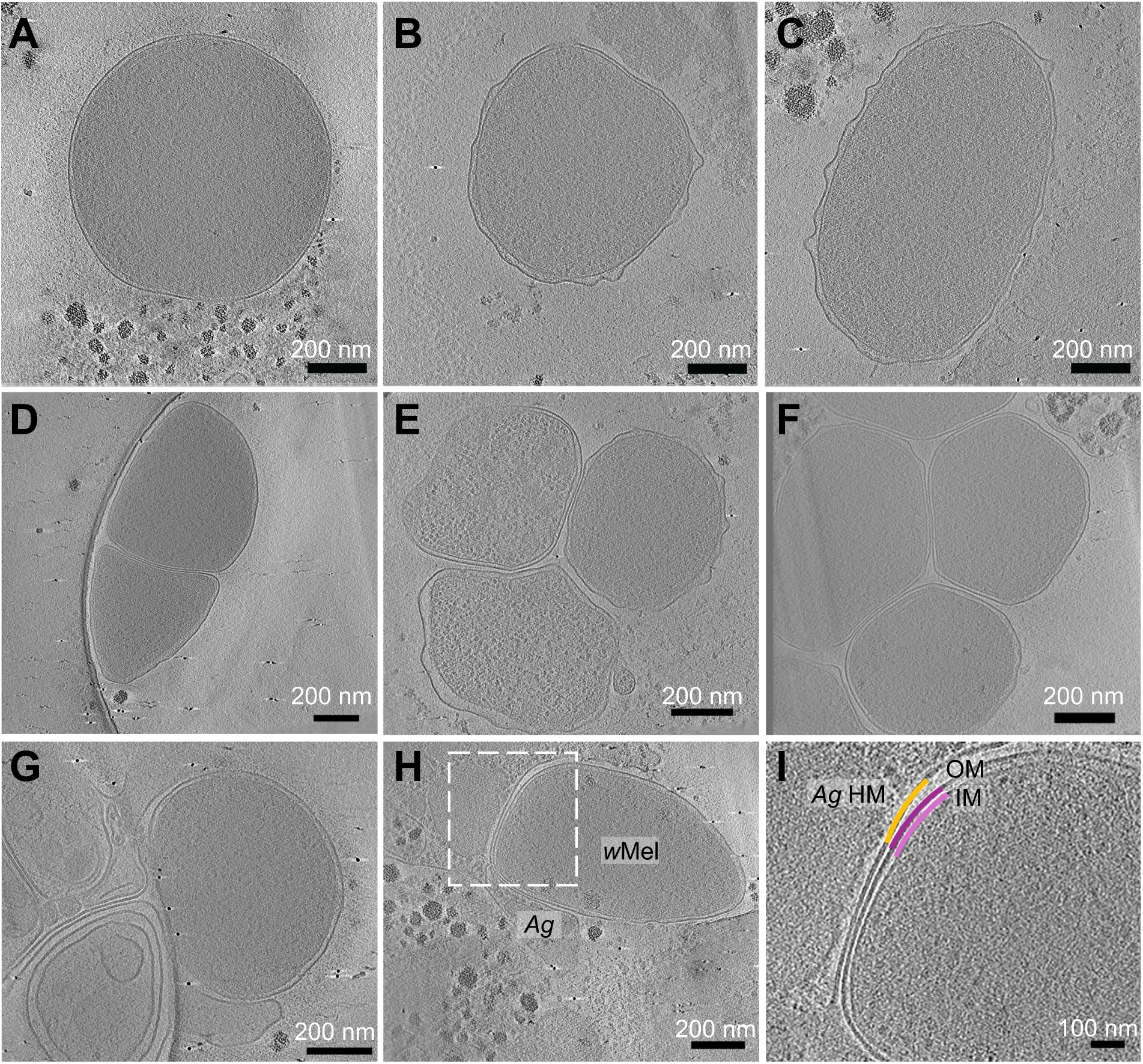
Morphology and ultrastructure of isolated *Wolbachia pipientis*. Tomographic slices from representative isolated *Wolbachia* cells exhibiting spherical coccoid (**A**), transitional coccobacillus (**B**), and elongated bacillus (**C**) morphologies. (**D**) Tomographic slice from a representative *Wolbachia* cell undergoing symmetric binary fission. Tomographic slices from representative clusters of isolated *Wolbachia* cells exhibiting ruffled (**E**) and smooth (**F**) outer membranes. In panel **E**, one cell is shedding a large membrane vesicle. (**G-H**) Tomographic slices through representative *Wolbachia* cells that remain associated with *Anopheles gambiae* host membranes and subcellular compartments. (**I**) Magnified view of the boxed region in **H** demonstrating the proximity of the *Wolbachia* OM to *A. gambiae* host membranes. Scale bars represent 200 nm in panels A-H, and 100 nm in panel I. OM, outer membrane; *w*Mel, *Wolbachia pipientis* str. *w*Mel; *Ag*, *Anopheles gambiae*; HM, host membrane.

### Identification of novel *Wolbachia* subcellular features

We next sought to identify *Wolbachia* structural components that enable persistent host cell parasitism and tissue colonization. Close inspection of our tomograms identified a periplasmic hat-like structure (**Fig. S1A-C**) with a similar morphology to analogous complexes observed across diverse bacterial phyla (*62*). Consistent with our previous observations (*62*), subtomogram averaging (**Fig. S1D**) revealed that the *Wolbachia* periplasmic hat-like structure is tethered to the IM with dimensions of 20 nm wide at the base (proximal to the IM) and exhibits two cytoplasmic densities spanning approximately ∼10 nm (**Fig. S1D**). In addition to ruffled and smooth conformations, the *Wolbachia* OM often formed tubular projections (**Fig. S1E,F**). Many cells also exhibited cytoplasmic membranous extensions that appeared as stacks or horseshoe-like vesicles (**Fig. S1G,H**) derived from IM invagination into the cytoplasm. We also observed dense clusters in the cytoplasm of many *Wolbachia* cells (**Fig. S1I-L**). These structures ranged from ∼10-20 nm in size and appeared as granular, amorphous densities randomly localized throughout the cytoplasm.

In some cells, we observed distinct ladder-like structures of approximately 10 nm wide and few hundred nanometers long (**Fig. 2, Fig. S1M-P,** and **Movie S2**) that frequently exhibited a bending pattern following the contour of the OM, suggesting structural flexibility as well as tethering along the length of the structure (**Fig. 2A, Fig. S2M-P,** and **Movie S2**). Ladders were typically associated with the *Wolbachia* cell surface (**Fig. 2** and **Fig. S1M-O**); however, we identified one example in which the ladder appeared to be dissociated from the *Wolbachia* OM (**Fig. S1P**). In one instance, a ladder appeared to be tethered to periplasmic basal body machinery that is potentially associated with apparatus assembly and biogenesis (**Fig. 2B**). Subtomogram averaging of ladder-like particles revealed distinct structural periodicity (**Fig. 2C**) with pairwise fibrillar subunits arranged in a striated, pyramidal conformation when viewed down the long axis (**Fig. 2D-E**). Isosurface renderings of ladder particle averages revealed vertebrae-like subassemblies with frayed edges and periodic cross-linking densities in direct contact with the *Wolbachia* outer membrane (**Fig. 2F-I**). In some tomograms, we resolved co-purified *Anopheles gambiae* actin filaments (**Fig. 2J**) interfaced with *Wolbachia* ladder particles on the bacterial cell surface (**Fig. 2I,J**), suggesting new hypotheses surrounding the nature and function of novel ladder structures. We speculate that analogous to rodlet filaments assembled on the *Streptomyces* spore coat (*63*), *Wolbachia* ladders may function as a specialized motility or anchoring mechanism that enables non-sessile endosymbiont translocation to specific host cell compartments during embryogenesis and somatic tissue colonization.

**Figure 2.**
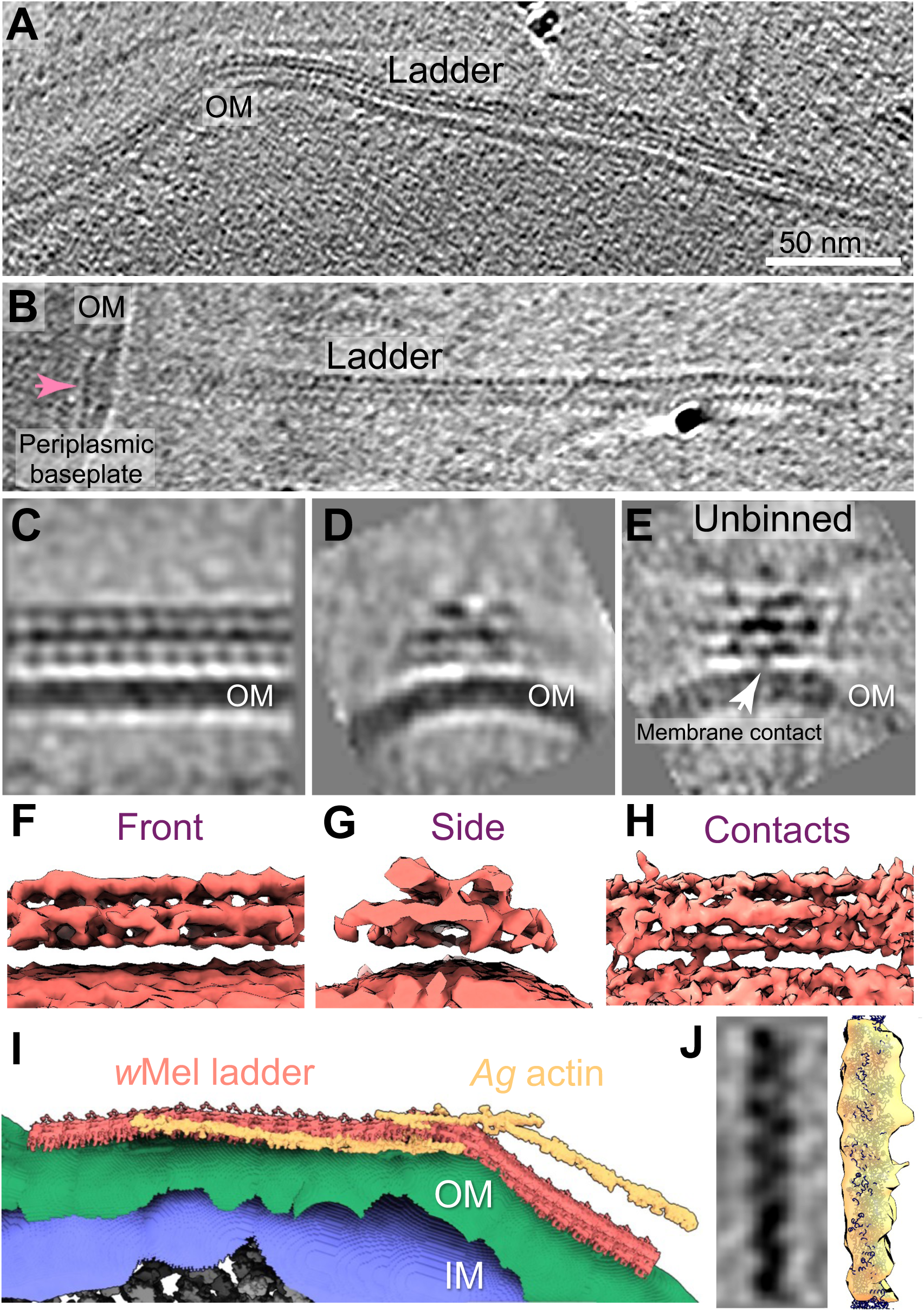
Structure of a ladder-like complex assembled on the *Wolbachia* cell surface. (**A**) Tomographic slice through a representative *Wolbachia* cell demonstrating the assembly of a ladder-like structure associated with the bacterial outer membrane. (**B**) Tomographic slice of a ladder structure exhibiting a periplasmic baseplate associated with apparatus biogenesis and/or tethering to the bacterial cell surface. (**C,D**) Subtomogram averages of ladder structures viewed *en face* (**C**) and down the long axis (**D**). (**E**) Unbinned subtomogram average of the ladder-like structure demonstrating membrane contacts with the bacterial OM. (**F-H**) Isosurfaces of the ladders depicted in **C-E**. (**I**) Segmentation of the *Wolbachia* cell depicted in A focused on the cell surface-associated ladder structure interfaced with co-purifying *Anopheles gambiae* actin filaments (yellow). (**J**) Subtomogram average (right) and segmentation (left) of *Anopheles gambiae* actin filaments. OM, outer membrane; IM, inner membrane; *w*Mel, *Wolbachia pipientis* str. *w*Mel; *Ag*, *Anopheles gambiae*. In **A**, scale bar represents 50 nm.

### *In vivo* ultrastructure of the α-proteobacterial *rvh* T4SS machinery

In some tomograms, we identified dense, disc-shaped particles spanning the bacterial periplasm that resembled the OM-associated core complex in previously characterized T4SS machineries (**Fig. 3**) (*64–67*). Based on structural similarities to pKM101 (*67*) and pED208 (*66*) conjugative systems, and to a lesser extent the *H. pylori cag* T4SS (*68, 69*) and *L. pneumophila dot/icm* T4SS (*70, 71*) complexes visualized *in situ*, we hypothesized that this structure corresponded to the *w*Mel *rvh* T4SS. Within individual *Wolbachia*, we observed various numbers of particles exhibiting multiple distinct layers of densities in the periplasmic space near the outer membrane and often arranged in arrays whereby multiple adjacent structures were assembled in close proximity (**Fig. 3B** and **Fig. 3C**, insets). Putative *w*Mel *rvh* T4SS complexes were localized to both the cell poles and lateral surfaces, consistent with *in situ* analysis of T4SS positioning in other Gram-negative organisms (*68, 70, 71*) (**Fig. 3**). Congruent with our hypothesis, many particles were associated with characteristic external conjugative pili (**Fig. 3**), leading us to conclude that this structure corresponds to the α-proteobacterial *w*Mel *rvh* T4SS nanomachine.

**Figure 3.**
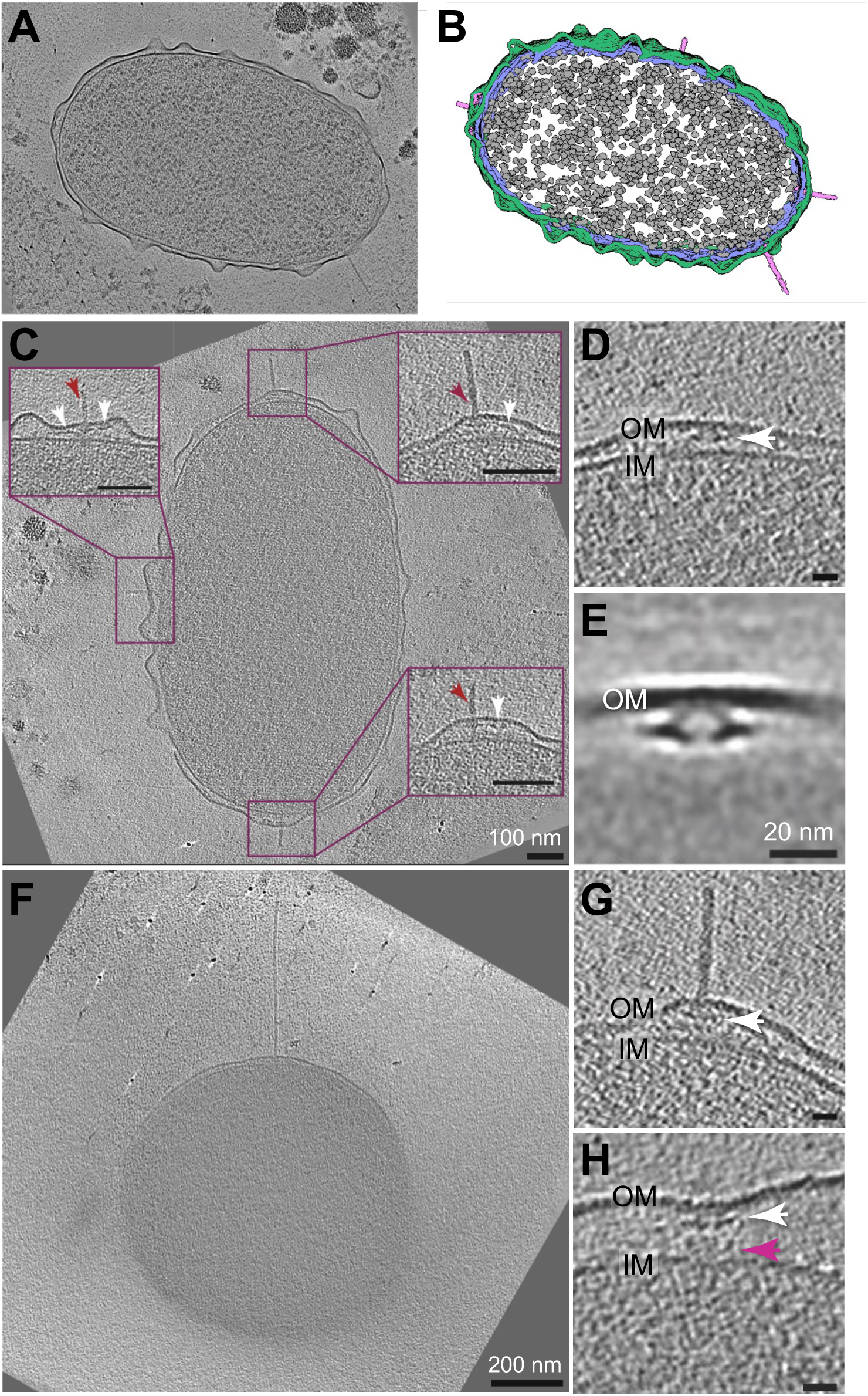
*In situ* architecture of the *w*Mel *rvh* T4SS apparatus. (**A**) Denoised tomographic slice and (**B**) segmentation of a representative bacillus *Wolbachia* cell exhibiting multiple piliated and non-piliated *rvh* T4SS nanomachines (pink) embedded between the bacterial outer membrane (green) and the inner membrane (blue). Ribosome densities in the cytoplasm are colored gray. (**C**) Tomographic slice through the cell depicted in **A** highlighting multiple *rvh* T4SS assemblies in the periplasm (magenta boxes). Piliated *rvh* T4SS machines (maroon arrows) are frequently positioned proximal to non-piliated *rvh* T4SS complexes (white arrows) organized in tandem “secretion arrays” (enlarged boxes). (**D**) Tomographic slice through a representative non-piliated *rvh* T4SS outer membrane complex particle (white arrow) associated with the cell envelope. (**E**) Central slice through the *rvh* T4SS outer membrane complex subtomogram average. (**F**) Tomographic slice through a representative spherical coccoid *Wolbachia* cell elaborating a rigid extracellular *rvh* T4SS pilus extending from the bacterial cell surface. (**G**) Tomographic slice through a piliated *rvh* T4SS particle (white arrow) exhibiting characteristic outer membrane complex structures and additional densities in the periplasm and inner membrane. (**H**) Tomographic slice through a fully assembled, non-piliated *rvh* T4SS particle encompassing an outer membrane complex (white arrow) and wing-like densities spanning the periplasm into the inner membrane (magenta arrow). OM, outer membrane; IM, inner membrane. In **D-E** and **G-H**, scale bars represent 20 nm.

In addition to piliated structures, we identified several non-piliated particle subassemblies within the periplasm. The most abundant T4SS particle form consisted of the OM-associated core complex (OMC) (**Fig. 3D**, white arrow). Positioned at the periplasmic face of the OM, the OMC adopted a disc-like structure and lacked obvious IM-associated components (**Fig. 3D**). We identified 72 OMC particles in 41 cells (**Table S1**), with significantly greater numbers visualized in *Wolbachia* with a ruffled OM (44 particles) versus a smooth OM (28 particles). To investigate structural details of the *w*Mel *rvh* T4SS, we generated a subtomogram average of the OMC (**Fig. 3E**). Subtomogram averaging of *w*Mel *rvh* T4SS OMC particles revealed a central “chamber” structure connected to the periplasmic face of the OM through weaker “arm” densities that appear to anchor the OMC chamber to the OM inner leaflet (**Fig. 3E**). Notably, in contrast to other T4SS architectures visualized *in situ*, the *w*Mel *rvh* T4SS OMC was associated with a disc-like periplasmic “baseplate” density (**Fig. 3E**). At the resolution of our cryo-tomograms, the overall *w*Mel *rvh* T4SS OMC architecture resembles the so-called OMC “flying saucer” densities visualized in other conjugative T4SS machineries (*66, 67*). However, in contrast to other T4SS architectures associated with OM distortion and remodeling required to accommodate OMC biogenesis (*66–71*), the *w*Mel *rvh* T4SS OMC was not associated with apparent OM deformation and we could not detect an obvious translocation pore (**Fig. 3D,E**).

The second most abundant form of *w*Mel *rvh* T4SS particles elaborated rigid, extracellular conjugative pili on the bacterial cell surface (**Fig. 3F,G**, and **Movie S3**). We identified 20 piliated T4SS particles that assembled rigid pili of various lengths ranging from a few nanometers to more than 500 nm (**Fig. 3** and **Movie S3**). Piliated *w*Mel *rvh* T4SS particles were frequently spatially clustered in arrays adjacent to non-piliated OMC subcomplexes (**Fig. 3B,C**) and were assembled by both bacillus (**Fig. 3A-C**) and spherical (**Fig. 3F**) *Wolbachia* morphologies. In addition to the non-piliated OMC, we identified non-piliated *w*Mel *rvh* T4SS particles comprising both the OMC, an associated thin “stalk” structure extending to the IM, and an IM-associated complex (designated the IMC) (**Fig. 3H**), albeit at a much lower frequency (only 4 such particles were identified, **Table S1**). In addition to the distinct densities in the OMC, *w*Mel *rvh* T4SS particles harboring the IMC were structurally reminiscent of diverse T4SS complexes observed *in situ*, including corresponding architectures of prototypical conjugative systems (*66, 67, 72*) and expanded machineries (*68, 70, 71*). In contrast to the pKM101 and pED208 conjugative systems (*66, 67, 72*), we did not observe *w*Mel *rvh* T4SS-associated pili on the *Wolbachia* cell surface in the absence of periplasmic basal body densities. We speculate that compared to paradigmatic conjugative pili, the *w*Mel *rvh* T4SS elaborates mechanically resilient pili that are not readily detached or shed from the cell surface. The observed *w*Mel *rvh* T4SS pili also appeared inflexible with blunt ends that cap an apparent lumen (**Fig. 4**), further demonstrating architectural relatedness to T4SS archetypes.

**Figure 4.**
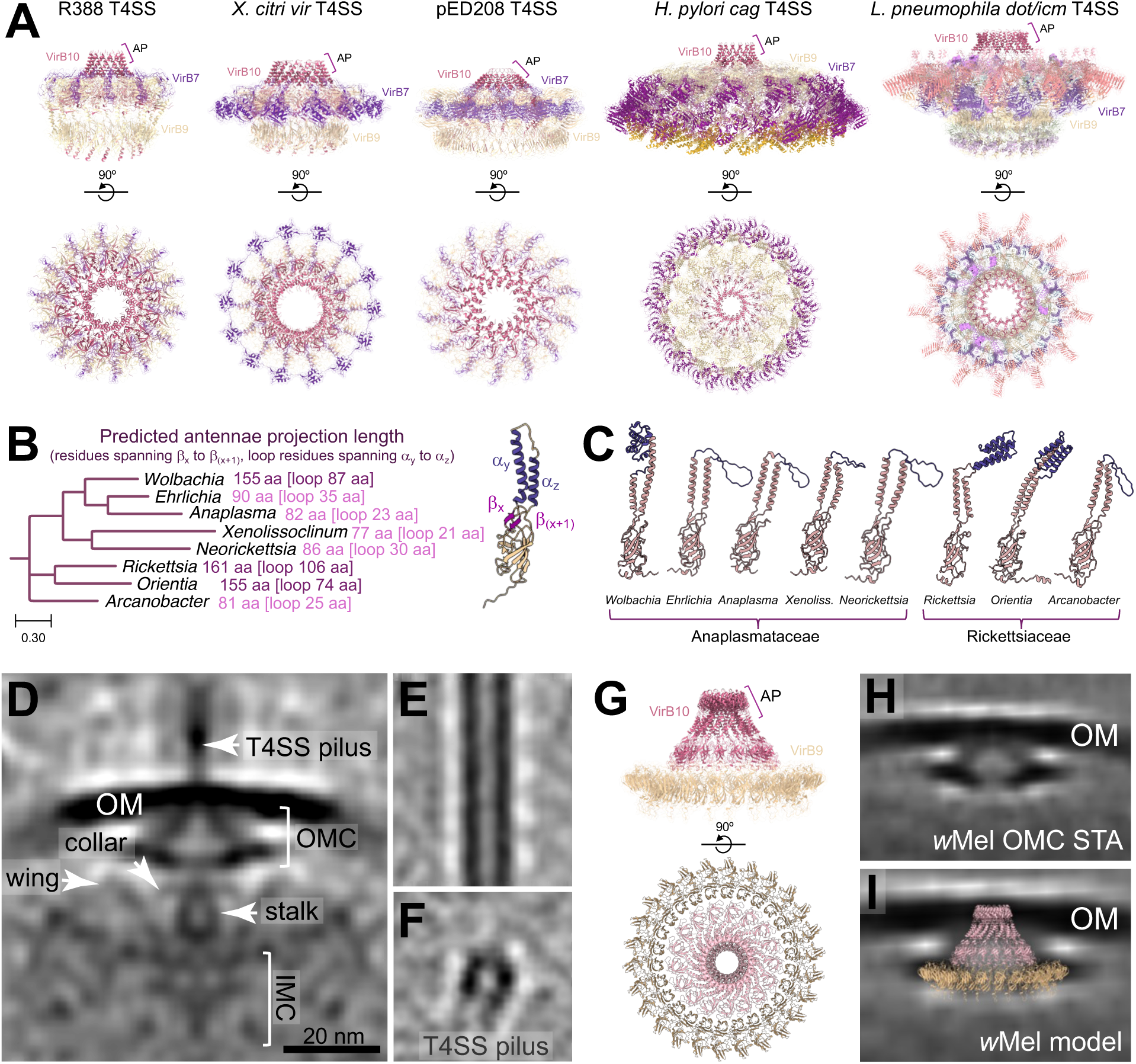
Structural modeling of the α-proteobacterial *rvh* T4SS nanomachine. (**A**) Molecular architecture of the T4SS outer membrane complexes assembled by diverse bacteria. The outer membrane spanning VirB10 antennae projections (AP) are a conserved feature among minimized (R388 T4SS and *X. citri vir* T4SS) and expanded (pED208 T4SS, *H. pylori cag* T4SS, and *L. pneumophila dot/icm* T4SS) machineries. Images depict the side view (top row) and top-down view (bottom row) of the single particle cryo-EM structures for each system [R388 plasmid-encoded T4SS (PDB 7O3J and 7O3T), *Xanthomonas citri vir* T4SS (PDB 6GYB), F plasmid-encoded T4SS (PDB 7SPC and 7SPB), *H. pylori cag* T4SS (PDB 6X6S), and *L. pneumophila dot/icm* T4SS (PDB 7MUY)]. (**B**) Phylogenomic reconstruction of selected Rickettsiales lineages (left). Scale bar indicates nucleotide substitutions per site. Numbers denote the predicted length of VirB10 antennae projections (spanning from β-strand β_x_ to β_(x+1)_) and flexible loop regions (traversing α-helices α_y_ to α_z_) for each predicted ortholog structure (aa, amino acids). Inset (right) depicts the structure of a prototypical VirB10 subunit from the R388-encoded conjugative T4SS (PDB 7O3J) demonstrating the positions of β-strands and α-helices that form the cognate antennae projection. (**C**) Atomic models of predicted VirB10 subunit C-terminal domains (pink) and associated antennae projections (purple) for the indicated species in the Rickettsiaceae and Anaplasmataceae lineages. In contrast to terminal, flexible loops in VirB10 orthologs harbored by related Rickettsiales lineages, *Wolbachia*, *Rickettsia*, and *Orientia* orthologs are predicted to exhibit expanded antennae projection subdomains composed of α-helix bundles extending from the top of conserved VirB10 β-barrel domains. (**D**) Central slice of the piliated T4SS nanomachine subtomogram average. Structural features associated with the piliated structure are highlighted. (**E,F**) Central slices through the side view (**E**) and cross-section (**F**) of the *w*Mel *rvh* T4SS pilus subtomogram average. (**G**) Structural modeling of the *w*Mel *rvh* T4SS outer membrane core complex demonstrating extended VirB10 antennae projections. (**H, I**) Docking the predicted *w*Mel *rvh* T4SS core complex models into the *in situ* cryo-ET *rvh* OMC density map (**H**) via rigid-body fitting (**I**) revealed a congruent integrative model in which the shape and size of the predicted structure docks into the appropriate densities. STA, subtomogram average; OM, outer membrane; OMC, outer membrane complex; IMC, inner membrane complex.

### Extended antennae projections in the *w*Mel *rvh* T4SS OMC

Conserved across all T4SS architectures, the OMC exhibits a channel-like, OM-spanning antennae projection (AP) subdomain formed by VirB10 α-helical elements connected by an unstructured loop (**Fig. 4A**) (*72–81*). Previous bioinformatic analyses of regions flanking predicted VirB10 APs revealed that AP length is conserved across Proteobacteria (with an average AP length of ∼57 residues) despite a lack of sequence conservation (*82*). However, the AP length of VirB10 orthologs in Rickettsiales lineages is predicted to be significantly longer (∼127 residues) than the corresponding regions of prototypical components (**Fig. 4B**), suggesting structural diversification of anomalous T4SS machineries harbored by obligate intracellular bacteria (*57, 82*). Compared to AP structures resolved in Proteobacterial T4SS complexes (**Fig. 4A**) in which the unstructured, flexible loop ranges from 16 to 32 residues in length (*79*), structural modeling of VirB10 orthologs in the Rickettsiales revealed striking diversity in AP loop length among Rickettsiaceae and Anaplasmataceae lineages (**Fig. 4C** and **Fig. S2A**). Consistent with previous bioinformatics analyses (*82*), *Rickettsia* (Rickettsiaceae), *Orientia* (Rickettsiaceae), and *Wolbachia* (Anaplasmataceae) exhibit exceptionally large putative AP loop insertions predicted to form structured two-helix bundles (**Fig. 4C** and **Fig. S2A**), while other rickettsial lineages retained unstructured loop regions with lengths similar to typical Proteobacterial VirB10 subunits (**Fig. 4C**). Coupled with phylogenomic analyses (*57, 82*), our studies suggest that extended AP loop structures evolved from ancestral VirB10 orthologs and were retained in diversified T4SS architectures harbored by specialized endosymbiont lineages.

To explore the structural relationships between the *w*Mel *rvh* T4SS and orthologous nanomachines, we next compared the piliated *w*Mel *rvh* T4SS subtomogram average to minimized [R388 (*73*) and *X. citri vir* T4SS (*74*)] and expanded [pED208 (*72, 83*), *H. pylori cag* T4SS (*76, 78*), and *L. pneumophila dot/icm* T4SS (*77*)] structures resolved by single particle cryo-electron microscopy (**Fig. 4A**) and *in situ* cryo-ET (**Fig. S3**). By averaging only the piliated tomograms, which contain fully assembled T4SS periplasmic complexes, we observed striking architectural similarity between the *Wolbachia rvh* T4SS complexes (**Fig. 4D-F**) and the expanded pED208-encoded T4SS machinery visualized *in situ* (**Fig. S3A**) (*66*) and in single particle cryo-EM studies (**Fig. 4A**) (*72*). Although the *w*Mel *rvh* T4SS is phylogenetically distinct from paradigmatic conjugation systems, subtomogram averaging of the assembled *w*Mel *rvh* T4SS revealed remarkable *in situ* structural resemblance to the pED208-encoded F pilus apparatus, including densities corresponding to the OMC, a periplasmic collar near the IM at the boundary of the IMC, a stalk-like cylinder extending from the OMC to the IMC, and weak cytoplasmic densities perpendicular to the IM resembling inverted “horseshoes” associated with the IMC (**Fig. 4D** and **Fig. S3A,B**). Conversely, prominent *w*Mel *rvh* OMC “baseplate” and “wing-like” densities were absent in the visualized pED208 architecture (*66*), revealing unique structural features in the *Wolbachia rvh* T4SS nanomachine (**Fig. 4D** and **Fig. S3A,B**). Consistent with previous structural studies of prototypical conjugative machineries (*64, 84–86*), subtomogram averaging of the *w*Mel *rvh* T4SS-associated conjugative pilus revealed a hollow lumen or channel exhibiting a diameter of approximately 10 nm (**Fig. 4E,F**).

In our subtomogram averages of non-piliated (**Fig. 3E**) and piliated (**Fig. 4D**) assemblies, we noted that the overall shape and diameter of the *w*Mel *rvh* OMC closely resembled the core complexes of both minimized and expanded T4SS nanomachine architectures assembled from VirB7, VirB9, and VirB10 subunits (*72–74, 83*). However, due to the low numbers of piliated *w*Mel *rvh* T4SS particles identified in our studies, the resolution of the subtomogram averages was insufficient to perform integrative structural analysis. Therefore, to gain insight into potential structural attributes associated with extended “baseplate” densities, we built a *w*Mel *rvh* OMC model using AlphaFold structural prediction (*87, 88*) and subsequent molecular docking to the pED208 core complex resolved by single particle analysis (**Fig. 4G** and **Fig. S2**). As with other Anaplasmataceae, *Wolbachia* exhibit *virB9* duplication and encode a single copy of *virB10* (*45, 46, 57*). Although a previous report identified a putative *virB7* ortholog in *Wolbachia w*Pip (WP0823) (*57*), we were unable to identify a corresponding VirB7 component in *w*Mel via structural homology-based approaches (e.g., FoldSeek (*89*)). Further, the putative *w*Pip VirB7, and other annotated VirB7 orthologs within the *Wolbachia* clade, are significantly truncated (74 residues) compared to the corresponding subunits in prototypical T4SS architectures. While we cannot rule out the possibility that the *w*Mel *rvh* T4SS incorporates truncated VirB7-like components into the OMC, we speculate that machinery diversification and extensive lineage-specific genome reduction in the Anaplasmataceae (*57, 82*) resulted in the exclusion of VirB7 orthologs in modern *w*Mel *rvh* T4SS architectures. We therefore omitted putative *w*Mel VirB7 orthologs from our OMC structural model.

Compared to the pED208 system, the *w*Mel VirB10 (WD_0006) C-terminal β-barrels assembled a predicted central cone subcomplex exhibiting extended α-helical AP subdomains predicted to span the OM (**Fig. 4G** and **Fig. S2**). Guided by our OMC subtomogram averages, we next modeled *w*Mel VirB9 (WD_0005) subunits into the pED208 structure. In contrast to the pED208 OMC which exhibits a peripheral ring composed of outer and inner layers assembled from VirB9, the predicted *w*Mel VirB9 model docked exclusively into the inner layer structure forming a ring complex with a similar shape and diameter as the disc-like “base plate” density (**Fig. 4G**). Superimposition of the *w*Mel *rvh* OMC subtomogram average (**Fig. 4H**) with either the pED208 central cone (**Fig. S2E**) or the *Wolbachia* VirB9/VirB10 structural model (**Fig. 4I**) revealed congruent density matching within the visualized subcomplexes. Consistent with structural diversification, the *w*Mel VirB9/VirB10 model exhibited an overall fit within the non-piliated (**Fig. 4I**) and piliated (**Fig. S3C**) *w*Mel *rvh* T4SS subtomogram averages, with the predicted VirB9 ring aligning to the OMC “baseplate” density and extended VirB10 APs positioned within the OM (**Fig. 4I** and **Fig. S3C**). Analysis of the pED208 (**Fig. S2F**) and *w*Mel (**Fig. S2G**) central cone hydrophobicity corroborated our imposition model and revealed a hydrophobic region near the OM insertion site, a hydrophilic chamber lumen, and hydrophilic AP α-helix bundles predicted to extend beyond the OM to potentially stabilize the assembled conjugative pilus or translocation channel pore (**Fig. S2E** and **Fig. S3C**). Finally, to identify architectural oscillations within the *w*Mel *rvh* T4SS basal body required to accommodate pilus biogenesis, we calculated difference maps between the piliated and non-piliated subtomogram averages **(Fig. S3D-F** and **Movie S4**). These analyses revealed several structural differences between the observed architectural states including (i) *rvh* T4SS pilus or stalk assemblies spanning the periplasmic apparatus, (ii) the positioning and conformation of putative VirB9 subassemblies in the *rvh* OMC “baseplate” structure, and (iii) the formation of stalk-like and inner membrane-associated densities in the piliated *rvh* T4SS average (**Fig. S3F** and **Movie S4**). Collectively, these studies provide the first visualization of α-proteobacterial *rvh* T4SS architecture *in situ* and reveal diverse structural innovations governing nanomachine biogenesis at the host-endosymbiont interface.

## DISCUSSION

Despite wielding a vast and significant influence on invertebrate biology, *Wolbachia* has never been visualized under native conditions at high macromolecular resolution, rendering many fundamental aspects of the endosymbiotic lifestyle hitherto unknown. Here, we applied cryo-ET to image *Wolbachia* cells purified from cognate insect cells, revealing many unexpected and novel structural features assembled by this fascinating symbiont. Similar to previous reports (*37, 41, 43*), we observed various *Wolbachia* morphologies ranging from bacillus and spherical forms to intermediate coccobacillary shapes. However, our studies uncovered additional insights into the morphological nature of the *Wolbachia* OM including both ruffled and smooth configurations (**Fig. 1**). We speculate that observed OM morphologies reflect various transitional states associated with distinct host or bacterial developmental stages. Moreover, we visualized the biogenesis of IM invaginations that create long membrane stacks in the cytoplasm. In other intracellular bacteria such as *Coxiella burnetii*, cytoplasmic stacks derived from the IM are hypothesized to coordinate morphological remodeling and rapid biphasic developmental transitions from small cell to large cell variants (*90, 91*). A similar hypothesis could explain our observation in *Wolbachia* whereby cytoplasmic IM invaginations could facilitate (or result from) the transition from one cell shape to another (**Fig. S1G,H**). Similarly, *Wolbachia* OM projections can harbor assembled protein subcomplexes (**Fig. S1E**), reminiscent of membranous tubes described in other species (*92*); however, whether OM-derived tubes or projections serve specific functional roles at the *Wolbachia*-host cell interface remains to be determined.

Our studies revealed other interesting structures positioned within the *Wolbachia* inner membrane including a “hat-like” structure (**Fig. S1A-D**) that we previously described in diverse bacterial species (*62*). At the resolution of the current tomograms, the general architecture and dimensions of the *Wolbachia* hat-like structure are similar to corresponding periplasmic densities observed in other species (*62*). However, this structure was previously linked to flagellar genes which are not encoded by *Wolbachia*. We speculate that the hat-like structure thus represents an ancient and broadly conserved machinery that assembles and functions independent of flagellar biogenesis. Our cryo-tomograms also revealed the presence of a cell surface-associated ladder-like structure tethered to the *Wolbachia* OM (**Fig. 2** and **Fig. S1M-P**). Reminiscent of rodlet pairwise aligned filaments assembled on the *Streptomyces* spore coat, we speculate that the *Wolbachia* ladder-like structure represents an analogous apparatus that facilitates cell translocation from the host cell cytoplasm. On a macro scale, *Streptomyces* rodlet layers provide a gripped surface that directly attaches to arthropods and nematodes for long range spore dispersal (*63*). On a micro scale, rodlet filaments comprise a cell surface mobilization platform whereby the spore becomes entangled and wrapped by flagella assembled by motile soil bacteria to mobilize non-sessile *Streptomyces* spores (*63*). In the case of *Wolbachia*, we propose that such a mechanism may enable immotile endosymbiont “hitchhiking” via direct association with actin (**Fig. 2I,J**) and microtubule networks to facilitate dissemination through diverse reproductive and somatic tissues. Given the importance of the actin cytoskeleton in host-endosymbiont interactions (*34, 35, 49, 51*), we hypothesize that the ladder-like structure is essential for *Wolbachia* germline transmission and cellular mobilization during dynamic intracellular lifestyles in somatic tissues.

Importantly, our work provides the first *in situ* structure of the α-proteobacterial *rvh* T4SS nanomachine (**Figs. 3-4**) and demonstrates that the *Wolbachia rvh* T4SS is morphologically similar to the pED208-encoded F pilus conjugation system at the macromolecular resolution of our cryo-tomograms (**Fig. S3**). In contrast to prototypical conjugative machineries that exhibit “arches” or “collars” associated with the inner membrane (*66, 67*), the *w*Mel *rvh* T4SS incorporates additional periplasmic “wing” densities similar to expanded effector-translocator architectures visualized *in situ* (**Fig. 4D** and **Fig. S3**) (*68–71*). We propose that the additional machinery decorations observed in the *w*Mel *rvh* T4SS arise from the numerous gene family expansions that underpin α-proteobacterial T4SS architectural complexity. For example, consistent with other Rickettsiaceae lineages, *Wolbachia* harbors multiple *rvh* T4SS gene proliferations including four tandemly arrayed copies of *virB6*, three *virB2* duplications, and multiple copies of *virB4*, *virB8*, and *virB9* scattered throughout the genome (*45, 46, 57, 82, 93*). Currently, the nature and function of *rvh* T4SS component duplications and expansions is a mystery. Rather than an artefact of evolutionary overengineering, we speculate that the α-proteobacterial *rvh* T4SS is an ideal example of a molecular Rube Goldberg contraption in which incorporation of diversified or duplicated components affords expanded subassembly and subcomplex complexity (*82, 94*). Such modular substructure assortments could thus facilitate context-dependent and specialized interactions with the host cell.

Curiously, in contrast to other visualized Proteobacterial T4SS architectures (*66–71*), the most frequently observed *w*Mel *rvh* T4SS complex comprised the assembled OMC tethered to the OM (**Fig. 3D**) without obvious periplasmic-spanning or inner membrane-associated structures. We speculate that analogous to other macromolecular machineries, including *Myxococcus xanthus* type IV pili (*95*) and the *Yersinia* type III secretion system (*96*), α-proteobacterial *rvh* T4SS apparatus biogenesis follows an outside-in assembly pattern whereby the OMC forms prior to the recruitment of periplasmic and IMC components. Alternatively, we cannot exclude the possibility that the observed *rvh* OMC subcomplexes represent T4SS particles that have undergone disassembly, leaving relic structures bound to the OM. Similar OM-associated disassembly relics resulting from dismantled flagellar motors have been previously described in multiple bacterial species (*97–103*). Future work to dissect the α-proteobacterial *rvh* T4SS apparatus assembly hierarchy in the context of host cell colonization, and the promise of future molecular biology approaches to manipulate the *Wolbachia* genome, will thus help to conclusively reveal the nature of the OMC subcomplex and *rvh* T4SS architecture observed here.

Paramount to our understanding of α-proteobacterial *rvh* T4SS assembly and function, and consistent with the observation that *Wolbachia* harbors three distinct *virB2* homologs, our cryo-tomograms definitively demonstrate that the *w*Mel *rvh* T4SS elaborates a long conjugative pilus capable of extending hundreds of nanometers beyond the bacterial cell surface. Currently, whether *w*Mel *rvh* T4SS pilus biogenesis is required for putative effector translocation into the target cell compartment is unknown and the mechanism by which *w*Mel VirB2 protomers polymerize during pilus extension in the absence of a *w*Mel VirB5 minor pilin/adhesin is unresolved. Furthermore, whether specific VirB2 subunits are functionally interchangeable or whether conserved pilin gene duplication reflects a redundant or specialized feature of the α-proteobacterial *rvh* T4SS machinery is enigmatic. One possibility is that specific VirB2 subunits are prioritized for extracellular pilus assembly while additional paralogs reinforce translocation channel integrity. Congruent with previous studies demonstrating that *virB2* paralog and putative effector gene expression positively correlates with (i) the L2 larval stage (*51*), (ii) pupation (coincident with the formation of reproductive tissues (*104, 105*)) (*45, 46, 51*), and (iii) in adult male flies (*45, 46*), we propose that the *w*Mel *rvh* T4SS conjugative pilus is assembled during discrete host developmental transitions to enable stage-specific and hierarchical effector translocation into the host cell. Thus, we hypothesize that microbial regulation of *w*Mel *rvh* T4SS pilus biogenesis imparts precise control of host manipulation and cellular interactions in response to conserved niche-specific developmental cues.

In addition to protein effector translocation, and given that *Wolbachia* genomic diversity is shaped by pervasive recombination and horizontal gene transfer (*106*), we speculate that the *w*Mel *rvh* T4SS pilus could facilitate interbacterial conjugative DNA delivery or alternatively mediate exogenous DNA import. Similarly, the observation that *Wolbachia* DNA can be horizontally transferred and integrated into cognate host genomes (*107–112*) raises the compelling hypothesis that the *w*Mel *rvh* T4SS pilus serves as a trans-kingdom conjugation conduit through which mobile genetic elements are transferred into the parasitized cell nucleus. While the current study provides the first visualization of α-proteobacterial *rvh* T4SS architecture, how the machinery is positioned within the bacterial envelope when interfaced with specific host cell compartments and whether the *rvh* T4SS-associated pilus penetrates host-derived vacuolar membranes is unknown. Future studies to resolve the *in situ* structure of the *w*Mel *rvh* T4SS apparatus assembled within the context of diverse host tissues and across the invertebrate developmental continuum from embryogenesis to adulthood will therefore provide mechanistic insight into the intricate nature and function of the specialized endosymbiont T4SS nanomachine.

Collectively, our work provides overarching insight into diverse cellular structures that orchestrate the dynamic choreography between endosymbiont and host. Furthermore, our studies provide the experimental framework for additional *in situ* cryo-electron tomography investigations to elucidate the structural basis of adaptations underlying the most prevalent intracellular infection on Earth. While the work here provides the first visualization of complex cellular architectures assembled by *Wolbachia*, several fundamental questions remain: What is the structural and mechanistic basis of cytoplasmic incompatibility and reproductive manipulation? How does *Wolbachia* colonization impart pathogen blocking? Does host range and interspecies engagement alter α-proteobacterial *rvh* T4SS machinery topology and architectural organization? Future *in vivo* structural biology studies performed in the context of host reproductive and somatic cell colonization will thus unravel multiple enigmatic aspects of the iconic *Wolbachia* endosymbiont to address long-standing questions in evolutionary biology, host-microbe symbiosis, speciation, and vector biocontrol.

## MATERIALS AND METHODS

### Cell lines and *Wolbachia* purification

*Wolbachia pipientis* strain *w*Mel was introduced into *Anopheles gambiae* Sua5B cells via direct transfer from infected *Drosophilae melanogaster* (*113*), and the symbionts were passaged in these established cells for several years prior to our experiments. For cryo-ET studies, *w*Mel was isolated from *Wolbachia*-infected *A. gambiae* Sua5B cells as previously described (*114*). Briefly, confluent *Wolbachia*-infected Sua5B cells were cultured in Schneider’s insect medium (Sigma-Aldrich) supplemented with 10% heat-inactivated fetal bovine serum (FBS) (Sigma-Aldrich). Monolayers were rinsed with sterile PBS and cells were detached mechanically using a sterile cell scraper. Cell pellets were obtained by centrifugation at 2,500 × g for 10 min at 4 °C and were resuspended in 10 ml fresh Schneider’s insect medium supplemented with FBS. Mechanical disruption was performed by vortexing the suspension for 5 min with sterile 3-mm borosilicate glass beads (∼100 per tube). The lysate was cleared by centrifugation at 2,500 × g for 10 min at 4 °C, and the supernatant was filtered through a 5-μm syringe filter (Millipore). *Wolbachia* were pelleted through a 250 mM sucrose cushion by centrifugation at 18,400 × g for 5 min at 4 °C. The final pellet was resuspended in 1 ml Schneider’s insect medium containing 10% FBS and further clarified by passage through a 2.7-μm syringe filter. *Wolbachia* density and viability were determined using the LIVE/DEAD BacLight Bacterial Viability Kit (Invitrogen) and quantified with a hemocytometer (*114*). Final bacterial preparations were adjusted to ∼10¹⁰ cells/ml.

### Sample preparation for cryo-electron tomography

Aliquots of isolated *Wolbachia* cells were mixed with fiduciary gold particles (Sigma Aldrich) that were concentrated and pre-treated with bovine serum albumin to prevent aggregation immediately prior to vitrification. Sample mixtures (3 µl) were deposited onto a freshly glow-discharged, R2/2 200 mesh Quantifoil EM grids (Quantifoil Micro Tools) and were plunge-frozen using a FEI Mark IV Vitrobot (Thermo Fisher Scientific, Waltham, MA). The Vitrobot chamber was set at 37°C with a relative humidity of 100%. The grid was blotted for 2.5 s and held for 1 s before vitrification by plunging into liquid ethane. Vitrified grids were stored under liquid nitrogen until loading into the microscope.

### Cryo-electron tomography data collection and processing

Tilt series were collected from −60° to +60° (at 1-2° tilt increments) at −8° µm defocus under low dose conditions on a Titan Krios G2 (Thermo Fisher Scientific) equipped with a K2 and Gatan Energy Filter at 4.3 Å/pixel. Exposure time was set to achieve a cumulative electron dose of ∼160 e^−^/Å^2^ for a complete tilt series. Tomograms were aligned and reconstructed using fiducial tracking in IMOD (https://bio3d.colorado.edu/imod/) (*115*) by weighted backprojection with contrast transfer function (CTF) correction. During reconstruction, tomograms were binned 3 times to 12.9 Å/pixel for increased contrast. For subtomogram averaging, particles were selected, aligned, and averaged in Dynamo (*116*). T4SS complexes were located by visual inspection, and dipole model points were placed at the center of each particle baseplate with north facing the outside of the cell. A default refinement within Dynamo was performed during alignment against an initial model generated from a subset of particles. In the first iterations, default spherical masks were used but a final round of alignments focused only on the periplasmic space was performed in all cases. Two-fold symmetry was applied to the *rvh* T4SS averages. For ladder complexes, the structure center lines were modeled as a filament in Dynamo and particle points were placed every 4 pixels with no rotational parameter before alignment by the default refinement settings against an initial model generated from a subset of particles. For F-actin averages, filament centerlines were modeled in Dynamo with a particle every 4 pixels before performing a global alignment. For T4SS structures, the following number of particles were used for subtomogram averaging: non-piliated *rvh* OMC, 70 particles; piliated *rvh* T4SS apparatus, 14 particles; *rvh* T4SS pilus, 73 particles. For the ladder-like structure, 253 particles from 5 individual structures were averaged. For the inner membrane-associated hat-like structure, 20 particles were averaged. For *Anopheles gambiae* F-actin, 141 particles were averaged.

Tomogram segmentations were performed in Dragonfly 2024 (https://dragonfly.comet.tech) (*117*). For segmentation, a 200 x 200 x 20 voxel slab was cropped from the edge of a single *Wolbachia* cell. The outer membrane, inner membrane, *rvh* T4SS particles, and ribosomes were hand segmented within the extracted slab, and the annotation was used to train a 5-layer 2.5D segmentation UNET. The parameters used for training were patch size: 128, stride ratio: 1, batch size: 16, loss function: categorical cross-entropy, augmentation factor: 5, and epochs: 300. Training was allowed to run until the loss value became asymptotic and was arrested due to an early stopping value of 15 epochs with no improvement. The trained network was inferred to the full tomograms, which were then hand-corrected if necessary. For the ladder-like structure, isosurface renderings of the subtomogram average were placed back into the segmentation and visualized using the ArtiaX plugin (*118*) for ChimeraX (*119*). Movies were subsequently scripted and rendered in ChimeraX. For visualizing the piliated *Wolbachia* cell depicted in Figure 3A and to aid in hand segmentation in Dragonfly, the tomogram was denoised with DeepClean, using our inhouse denoising network trained with outputs from CryoTomoSim (CTS) as previously described (*120*). Briefly, “synthetic cytoplasm” was simulated from a dense mixture of multiple, randomly oriented PDB files of varying size and shape and simulated membrane vesicles. Next, both a noisy and noise-free “prior” were simulated and used to train a regression-based UNET in Dragonfly (*117*). This network was then inferred to the *Wolbachia* tomogram.

### Phylogenetic tree construction

Bioinformatic analyses and phylogenetic reconstructions were made using pipelines available at the Bacterial and Viral Bioinformatics Resource Center (BV-BRC) (*121*). For whole genome phylogenetic tree analyses, the BV-BRC Codon Tree method was used to select individual, single-copy PAThosystems Resource Integration Center (PATRIC) global protein families (PGFams) and trees were built from selected genomes by aligning proteins (and aligned with MUSCLE v5 (*122*)) and coding DNA from single-copy genes using the program RAxML (*123*) with 100 rounds of fast bootstrapping and default parameters. The following representative strains were included in the final whole genome phylogenetic tree: *Neorickettsia risticii* str. Illinois, *Candidatus* Xenolissoclinum pacificiensis L6, *Anaplasma centrale* str. Israel, *Ehrlichia chaffeensis* str. Liberty, *Wolbachia pipientis* strain *w*Mel, *Rickettsia typhi* str. TH1527, *Orientia tsutsugamushi* str. Boryong, and *Candidatus* Arcanobacter lacustris str. SCGC AAA041-L04. The final Newick file was rendered in TreeViewer version 2.2.0 (*124*).

### Structural Modeling

Atomic models of T4SS components were predicted using AlphaFold/ColabFold (*87, 88, 125*). Predicted VirB10 sequences were obtained from UniProt for the following representative bacterial strains: *Rickettsia typhi* str. Wilmington [Q68X80], *Orientia tsutsugamushi* str. Boryong [A5CBW7], *Neorickettsia sennetsu* str. Miyayama [Q2GD29], *Anaplasma phagocytophilum* [Q8RPL5], *Ehrlichia chaffeensis* [Q8RPM1], *Wolbachia pipientis* str. *w*Mel [Q73IY6], *Arcanobacter lacustris* [A0A0F5MPH3], and *Xenolissoclinum pacificiensis* L6 [W2V115]. VirB10 sequences were aligned and trimmed to the pED208 T4SS OMCC central cone (PDB 7SPC) and atomic models were visualized in ChimeraX (*119*). For *Wolbachia rvh* T4SS OMC modeling, predicted VirB10 (WD_0006, UniProt Q73IY6) and VirB9 (WD_0005, UniProt Q73IY7) structures were visualized in ChimeraX and were aligned to pED208 T4SS scaffolds (PDB 7SPC and 7SPB) using the *matchmaker* command. T4SS models were docked into subtomogram averages in ChimeraX using the *fitmap* command and manual alignment into the appropriate densities.

## ACKNOWLEDGMENTS

The authors thank Dr. Susan Hafenstein (University of Minnesota) for helpful discussions. This study was conducted with federal funds from the National Institute of Allergy and Infectious Diseases (R01AI116636 to J.L.R), the National Institute of General Medical Sciences (P20GM130456 to C.L.S. and R35GM157116 to M.K.), the Searle Scholars Program (to M.K.), and seed funding from the Huck Institutes of the Life Sciences, Pennsylvania State University (to S.P.). J.L.R. was additionally supported by USDA Hatch project 4769, and funds from the Dorothy Foehr Huck and J. Lloyd Huck endowment. The Pennsylvania State University cryo-EM and cryo-ET Core (RRID:SCR_021178) services and instruments used in this project were funded, in part, by the Pennsylvania State University College of Medicine via the Office of the Vice Dean of Research and Graduate Students and the Pennsylvania Department of Health using Tobacco Settlement Funds (CURE). The content of this manuscript is solely the responsibility of the authors and does not necessarily represent the official views of the National Institutes of Health or the Pennsylvania State University. The Pennsylvania Department of Health specifically disclaims responsibility for any analyses, interpretations, or conclusions.

## AUTHOR CONTRIBUTIONS

Conceptualization: S.P., J.L.R., and M.T.S. Investigation: S.P. purified and prepared *Wolbachia* for imaging studies. J.H. and M.T.S. performed cryo-ET sample preparation, cryo-ET imaging, and data reconstructions. C.L.S. performed integrative structural modeling and bioinformatics studies. M.T.S., J.H., C.L.S, E.R., T.Z., M.K. performed data reconstruction and analysis. Methodology: M.T.S. designed data analysis and imaging methods. Formal Analysis: M.T.S., C.L.S., and M.K. analyzed, processed, and interpreted the cryo-ET data. Resources: M.T.S. provided cryo-ET resources. J.L.R. provided cell lines for *Wolbachia* purification. Writing – original draft: C.L.S. and M.K. Writing – review and editing: S.P., J.L.R., M.T.S., C.L.S., and M.K. All authors reviewed and approved the final manuscript draft. Visualization: M.T.S., C.L.S., and M.K. Supervision: M.T.S., C.L.S., M.K, and J.L.R. Funding acquisition: S.P., C.L.S., M.K., and J.L.R.

## DATA AVAILABILITY

Project data are deposited and available in the Dryad Digital Repository at doi.XXXXYYY.

---

**Figure S1.**
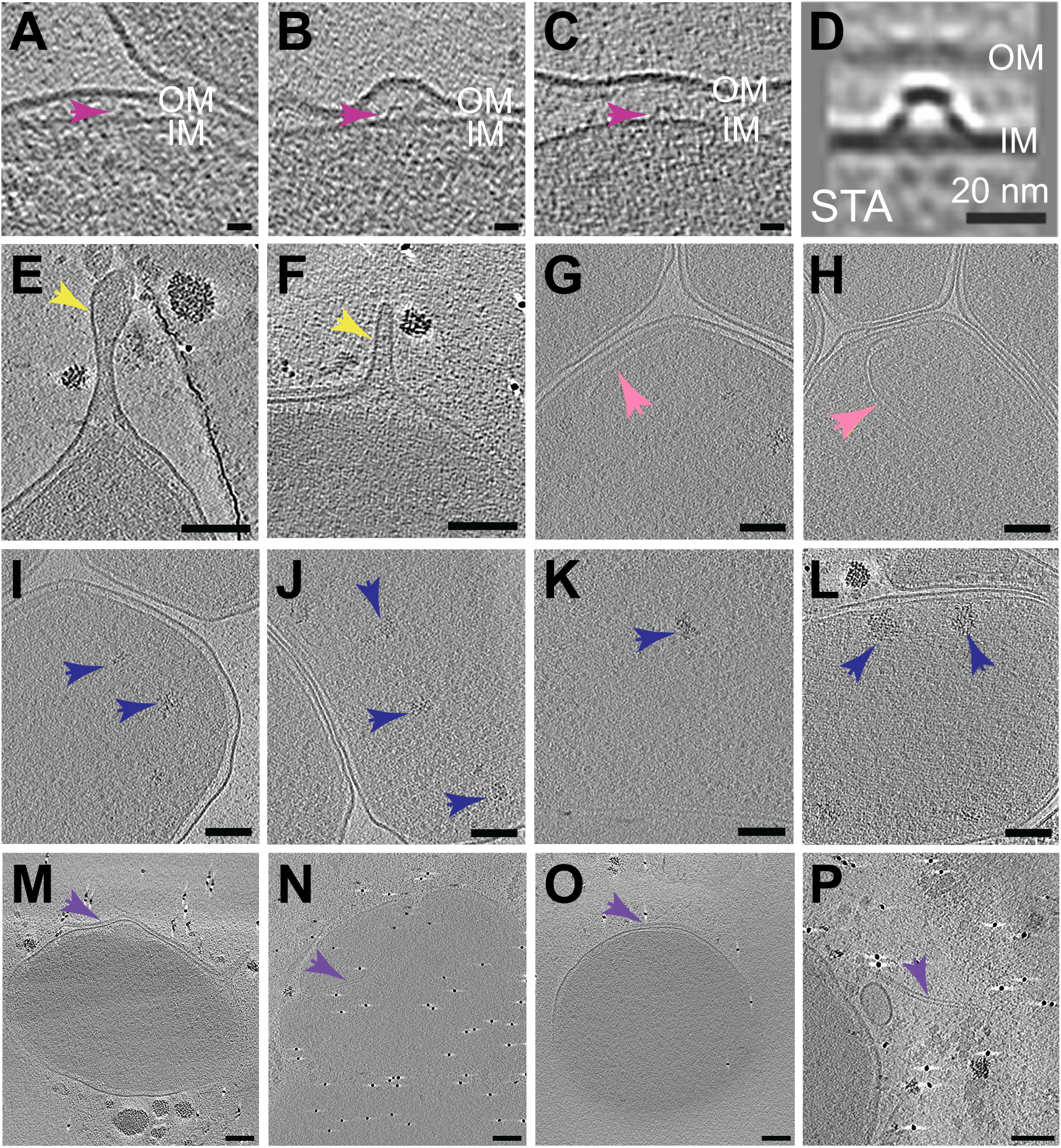
*In situ* visualization of *Wolbachia* cellular structures and macromolecular complexes. (**A-C**) Tomographic slices through representative *Wolbachia* cells exhibiting a periplasmic hat-like particle (magenta arrow). (**D**) Subtomogram average of the *Wolbachia* hat-like particle. (**E-F**) Tomographic slices through representative *Wolbachia* cells with outer membrane-derived projections (yellow arrows). (**G-H**) Tomographic slices through representative *Wolbachia* cells with inner membrane invaginations (pink arrows). (**I-L**) Tomographic slices through representative *Wolbachia* cells with amorphous, cluster-like densities in the cytoplasm (blue arrows). (**M-P**) Tomographic slices through representative *Wolbachia* exhibiting ladder-like structures associated with the outer membrane (purple arrows). In **P**, the ladder structure appears to be disassociated from the bacterial membrane. OM, outer membrane; IM, inner membrane. In all panels, scale bars represent 20 nm.

**Figure S2.**
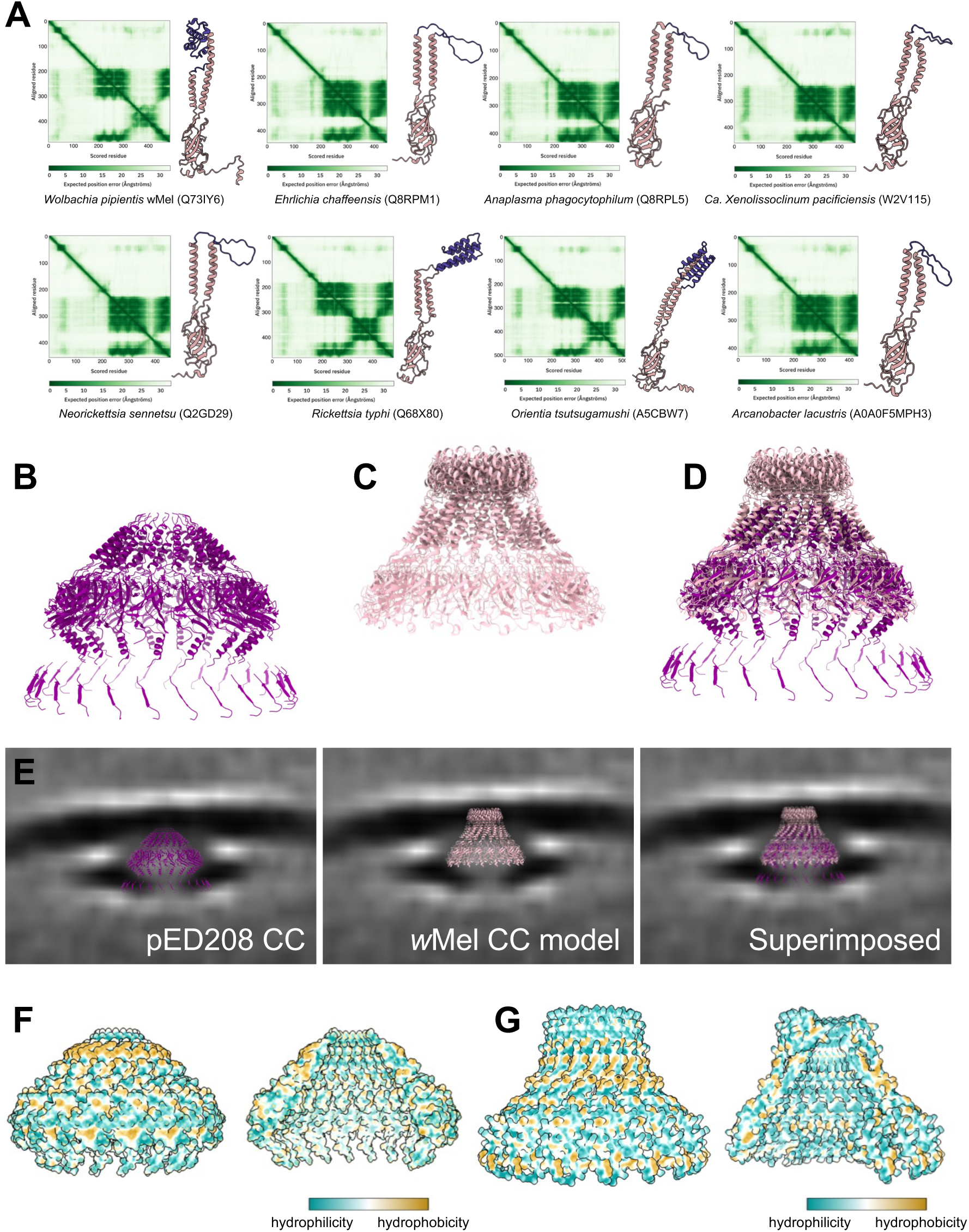
Antennae projection structural modeling and subunit conservation in diverse T4SS architectures. (**A**) AlphaFold models of Rickettsiales VirB10 subunit C-terminal domains. Predicted Aligned Error (PAE) plots indicate alignment error (in Å) for a pair of residues, indicating the positional uncertainty at residue x if the predicted and actual structures are aligned on residue y. For all predicted structures, the VirB10 C-terminal domain exhibits structural conservation with characteristic β-barrel domains and α-helical antennae projections. UniProt entries are indicated in parentheses. (**B**) Ribbon diagrams of the pED208-encoded T4SS central cone and (**C**) predicted *w*Mel VirB10 core complex. (**D**) Superimposition of pED208 central cone subassembly and the predicted *w*Mel VirB10 core complex demonstrating structural conservation. (**E**) Rigid-body fitting of the pED208-encoded central cone (left), the predicted *w*Mel *rhv* T4SS central cone (center), and superimposed core complex structures (right) in the *w*Mel T4SS OMC *in situ* cryo-ET map. Docking supports the assignment of the visualized densities as the putative α-proteobacterial *rvh* T4SS. (**F-G**) Hydrophobicity analysis of VirB10 antennae projection subdomains in the pED208-encoded T4SS central cone (**F**) and the *w*Mel *rvh* T4SS OMC (**G**). Molecular hydrophobicity potential demonstrates hydrophobic regions (yellow) near the outer membrane insertion point, with hydrophilic chamber lumens (cross-sectional views) and antennae projections (blue). OM, outer membrane; CC, central cone.

**Figure S3.**
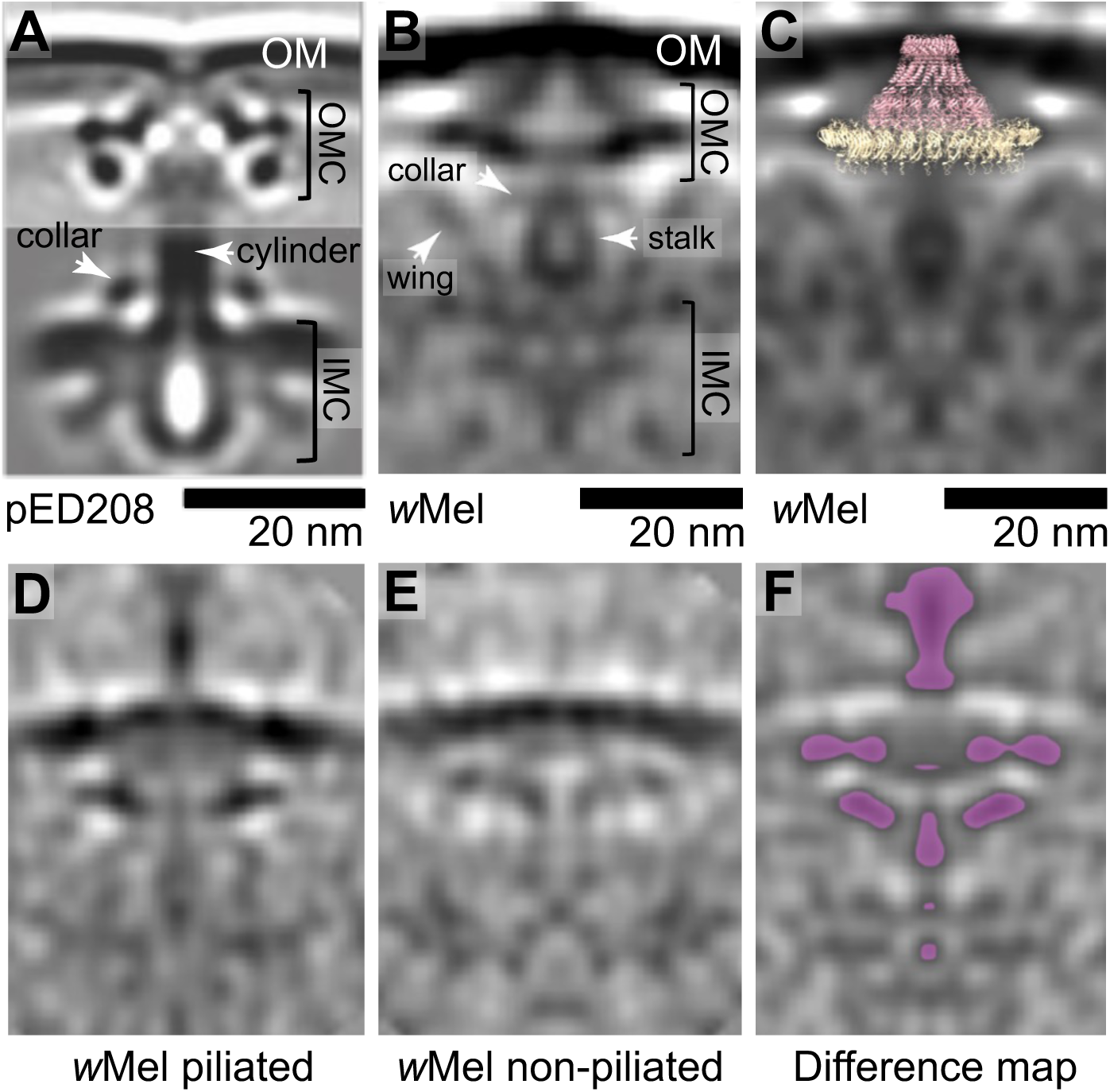
Architectural comparison of the pED208-encoded and *w*Mel *rvh* T4SS machineries. (**A**) Central slice through the *in situ* cryo-ET subtomogram average of the pED208-encoded T4SS (EMD-9344). (**B**) Central slice through the piliated *w*Mel *rvh* T4SS subtomogram average highlighting architectural features. (**C**) Rigid-body docking of the predicted *w*Mel *rvh* T4SS OMC in a tomographic slice of the piliated *rvh* T4SS cryo-ET map demonstrating shape and size similarity between the assembled OMC densities and the modeled *w*Mel VirB9/VirB10 subcomplex. (**D-F**) Central tomographic slices through the density maps comparing the piliated (left) and non-piliated (center) *rvh* T4SS particles. Difference map (right) indicates intensities corresponding to density differences (lavender) overlaid on the non-piliated particle. OM, outer membrane; OMC, outer membrane complex; IMC, inner membrane complex.

**Table S1:** *Wolbachia* cellular features described in this study.

| Feature | Number of observations |
| --- | --- |
| Total number of *Wolbachia* cells | 41 |
| Bacillus cell morphology | 11 |
| Spherical/coccoid cell morphology | 15 |
| Intermediate cell morphology | 15 |
| wMel *rvh* T4SS OMC (non-piliated) | 72 |
| Non-piliated wMel *rvh* T4SS (OMC and IMC) | 4 |
| Piliated wMel *rvh* T4SS | 20 |
| Ladder-like structures | 6 |
| Periplasmic hat-like structure | 20 |
| OM tubular extensions | 2 |
| Cells with smooth OM | 23 |
| Cells with ruffled OM | 18 |
| Cytoplasmic amorphous cluster densities | 10 |
| Cytoplasmic membranous structures | 6 |

